# One tree, many colonies: colony structure, breeding system and colonization events of host trees in tunneling *Melissotarsus* ants

**DOI:** 10.1101/2020.10.21.348797

**Authors:** Pierre-André Eyer, Edward L. Vargo, Christian Peeters

## Abstract

Ants exhibit a striking variety of lifestyles, including highly specialist or mutualist species. The minute blind workers of the African genus *Melissotarsus* chew tunnels in live trees to accommodate their obligate partner scale insects. Their modified legs are adapted for tunneling, but are unsuited for walking outside, confining these ants to their initial host tree. Here, we investigated whether this unique lifestyle results in complex patterns of genetic diversity at different scales, from the same tree to different populations. Using 19 microsatellite markers, we assessed their mating strategy and colony structure among and across populations in South Africa. We showed that only one queen reproduces within a colony, mated with up to three males. Yet, several inseminated dealate queens are present in colonies; one probably replaces the older queen as colonies age. The reproduction of a single queen per colony at a given time results in genetic differentiation between colonies, even those located on the same tree. Overall, we discussed how the slow process of colony digging under the bark and the lack of worker patrolling above the bark might result in reduced competition between colonies and allow several *secluded* colonies to cohabit in a cramped space on a tree.

## Introduction

Ants are one of the most ecologically dominant insects due to a striking variety of diets (live and dead insects, fungi, honeydew and other sweet secretions), as well as a huge biomass resulting from large numbers of foraging workers that sustain their perennial colonies. Among 13,000 known species with colony sizes ranging over six orders of magnitude (<10 to millions of workers), some genera stand out for their extreme lifestyles, such as army-ants, social parasites, fungus farmers, and intimate mutualists with plants, sap-feeding insects or other organisms (Holldobler & Wilson 1990; Dill et al. 2002; Brady et al. 2006; Oliver et al. 2008). In addition to having profound ecological consequences, these extreme lifestyles can have major influences on the population genetic structure of the species. This is especially true for specialist or mutualist species because their dispersal abilities, hence the amount of gene flow within and between populations, are intrinsically linked to the local distribution of the partner organism (Herre et al. 1994; Hoeksema & Bruna 2000; Thompson 1999; Thompson & Cunningham 2002). Consequently, disruptive effects of geographical distance and/or barriers may be amplified in specialist and mutualist species by the patchy distribution of suitable habitats.

In addition to extreme lifestyles, ants are also characterized by a large diversity of mating strategies and colony composition, resulting in a range of genetic structure patterns. The number of reproductive queens in a colony varies greatly, together with the number of matings for each queen. The level of polygyny (multiple queens per colony) may also vary across populations within a single species (Ross & Keller 1995; Seppa et al. 2004; Purcell et al. 2015), as well as the degree of polyandry (multiple matings per queen) to a lesser extent (Boomsma & Van Der Have 1998). These strategies strongly affect dispersal and therefore differentially influence gene flow across populations, as monogyne and polygyne colonies frequently exhibit different modes of dispersal. Polygyny is often associated with dependent colony foundation (DCF), whereby new queens disperse on foot with nestmate workers to establish a new colony nearby (Cronin et al. 2013). Such short-range dispersal reduces the level of gene flow and often leads to increased genetic differentiation between populations (Liautard & Keller 2001; Clemencet et al. 2005; Leppanen et al. 2013). In contrast, monogyny is often associated with independent colony foundation (ICF), whereby new queens disperse through either nuptial flights or female-calling (Cronin et al. 2013, Peeters & Aron 2017). This long-range dispersal increases gene flow and usually decreases population structure. In addition to distinct mating strategies, ants also exhibit different colony structures, ranging from monodomy whereby each colony is made of a single nest to polydomy where colonies comprise several nests exchanging workers, brood and reproductive queens (Chapuisat et al. 2005; Jackson et al. 2007; Steiner et al. 2007; Helantera et al. 2009; Eyer et al. 2018a).

The African genus *Melissotarsus* can be included in this compendium of extreme lifestyles as it combines a number of unorthodox traits: these minute blind ants (2 mm) chew tunnels in live trees in order to accommodate diaspidid scale insects that are their obligate partners (Peeters et al. 2017 and references therein). Adaptations of the workers for tunneling under the bark include highly modified back legs that are incompatible with walking outside host trees (Khalife et al. 2018). Consequently, these highly specialized ants depend entirely on the diaspidids for food, even though these produce no honeydew (Peeters et al. 2017). Furthermore, adult queens and workers produce silk (unique among Formicidae) to secure their tunnels against arboreal ants (Billen & Peeters 2020), a crucial adaptation because their sting has been lost. In contrast, queens can fly and retain normal leg morphology, which suggests independent foundation of new colonies (ICF).

Because *Melissotarsus* workers cannot walk outside their tunnels, DCF can be ruled out. Colonies are therefore confined to the initial host trees and cannot relocate or expand to another tree. Because workers cannot chew tunnels in branches lacking sufficient bark thickness, colony foundation is restricted to established trees. Several incipient colonies may consequently inhabit the same tree. As the different colonies grow larger over the years, workers extend the network of tunnels throughout the host tree up to the highest branches, concurrently with tree growth. Twenty-three botanical families of trees have been recorded with *Melissotarsus* in Africa (Peeters et al. 2017), and these exhibit a substantial diversity of growth forms. In *M. beccarii* and *M. weissi* in Cameroon, behavioral observations suggested that the absence of aggression between different colonies may lead to colony merging, resulting in a single huge polygyne colony covering an entire tree (Mony et al. 2007). This hypothesis is based on conjecture and it remains unclear whether expanding colonies can mix freely within a tree, or whether strict colonial boundaries are maintained. Similarly, the cryptic life style of *Melissotarsus* hampers the recognition of colony boundaries, estimation of queen number per colony, whether all queens reproduce after putative merging of colonies, and whether additional queens are recruited as colonies age. If the latter, are these queens related to the founding queen? Or are they unrelated queens coming from other colonies? Likewise, several dealate queens are mated in a colony (Mony et al. 2002; Peeters et al. 2017), and it is unclear whether these queens mated with their brothers as the colony grows.

In this study, we investigated mating strategy and colony structure of the wood-chewing *Melissotarsus* ants across five populations in southern Africa. We questioned whether their confined foraging strategy results in complex genetic patterns among and between colonies. We sampled 35 nests of *Melissotarsus* in four localities in South Africa and one locality in Mozambique, and genotyped their workers at 19 microsatellite markers. We assessed the reproductive system and population genetic structure by exploring patterns of genetic diversity at different scales. We investigated genetic differentiation from the local scale (*i.e.,* nests located on different parts of same branch and nests located on different stems of the same tree) to the regional scale (*i.e.,* populations). We inferred the social structure and breeding system of each colony, determining the number of queens, the number of matings per queen and their mode of dispersal.

## Materials & Methods

### Colony sampling

*Melissotarsus beccarii* was studied in four locations across South Africa (2017-2019): uMkhuze (MK), St Lucia (SL), Eastern Cape (EC) and Cederberg (CE) (Figure 1; detailed sampling is provided in Supporting Information Table S1). Nests were found in different host trees according to location. However, the same diaspine species, *Morganella conspicua,* was identified in almost all the nests sampled. Inhabited trees were identified by veinlike markings on the bark, revealing the presence of tunnels under the surface. For each nest, small areas of bark were shaved off and at least 20 adult workers and larvae were collected in ethanol for genetic analyses. All trees sampled were mapped and documented with Mapit GIS.

**Figure 1:**
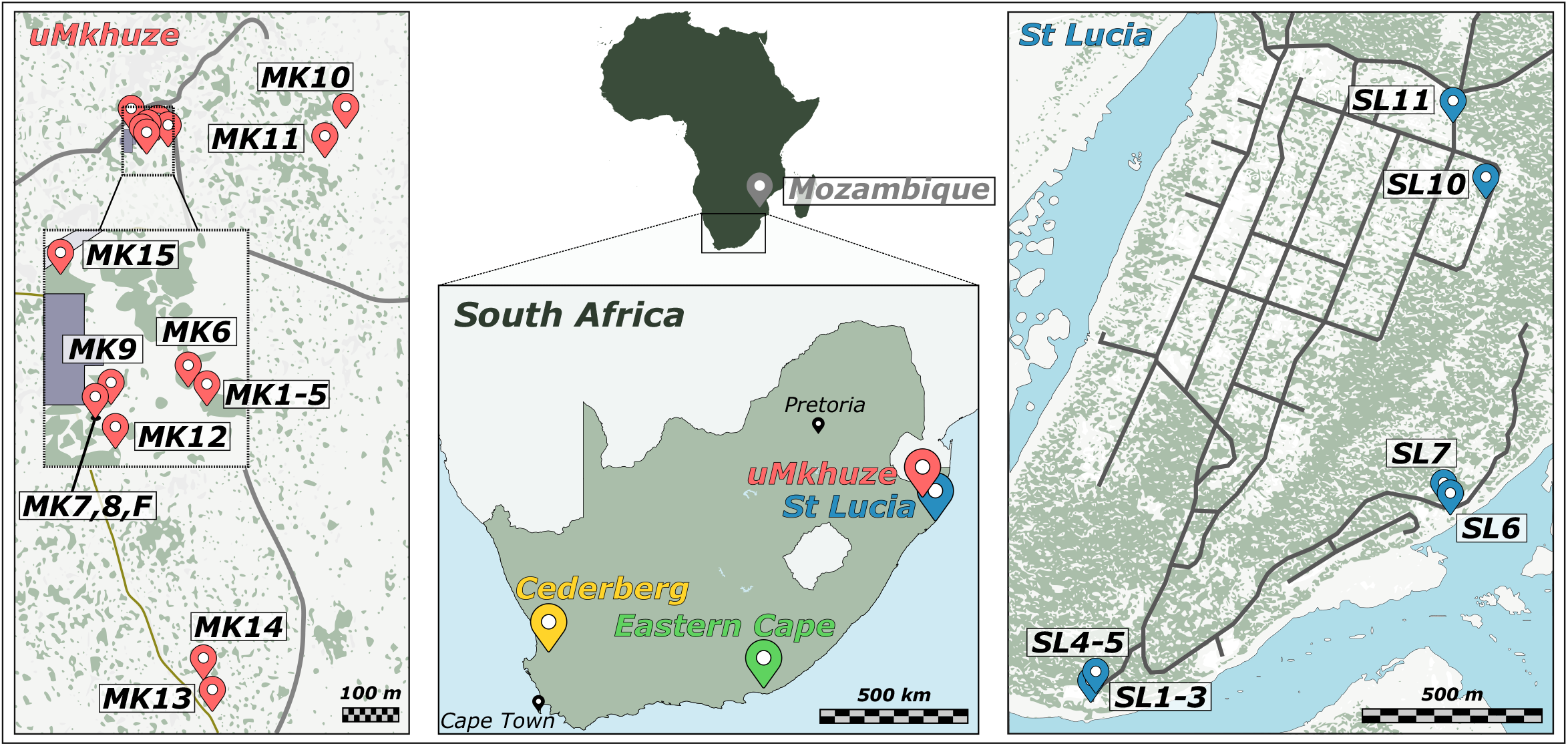
Locations of the 34 nests of *Melissotarsus* sampled in four localities in South Africa, and one pooled sample from Mozambique. Insets indicate sampling locations of nests within the populations of uMkhuze (left) and St Lucia (right). Nests located on the same branch or tree are indicated with the same label.

The uMkhuze and St Lucia locations in KwaZulu-Natal province are about 100km distant from each other. In uMkhuze, 17 nests were sampled, including six nests located on six different stems of the same *Strychnos madagascariensis* tree (*MK1*-*MK5*), and three nests located on different parts of the same stem (*MK7* + *MK8* + *MKF*; Figure 1; Supporting Information S1). *MKF* denotes an incipient founding colony made of three adults and five larvae. At this location, all but one nest inhabited *Strychnos madagascariensis* established trees (*MK15* was collected in *Ziziphus mucronata*). For four trees, sections of branches (50–60 cm long) were sawed off and taken to Paris (iEES) for behavioral observations and to search for reproductive queens. In St Lucia, nine nests were sampled out of five *Erythrina lysistemon* trees, including some nests (*SL1-3* and *SL4-5*) located on different stems of the same tree. For two trees, a branch was sawed and brought to Paris (Figure 1; Supporting Information S1). In the Eastern Cape population, four nests were sampled from *Olea capensis* and *Pterocelastrus tricuspidatus* trees. In the Cederberg (Western Cape), four nests were sampled from *Leucospermum praemorsum* and *Maytenus oleiodes*. Additional samples were collected in Mozambique (2016): Cabo Delgado province in the north (Namoto Forest; *Olax dissitiflora*) and Sofala province (Gorongosa National Park; *Piliostigma thoningii*). However, the Mozambique workers had been pooled and therefore were not used to characterize mating system and colony structure.

### Molecular analyses

DNA was extracted for eight randomly chosen workers per nest, all mother queens available and up to 8 alate queens per nest, following a modified Gentra-PureGene protocol (Gentra Systems, Inc. Minneapolis, MN, USA). Every individual was genotyped at 19 microsatellite markers previously developed for all species of ants (Butler et al. 2014; Supporting Information Table S2). Genotyping was performed using the M13-tailed primer method (Boutin-Ganache *et al.* 2001), which separately dyes each marker with a 5’-fluorescently labeled tail (6-FAM, VIC, PET or NED dyes). PCR reactions were carried out in a volume of 15 μL including 0.25-1.0 U of MyTaq™ HS DNA polymerase (Bioline), 2 μL of MyTaq™ 5x reaction buffer (Bioline), 0.08 μL of each primers, 0.08 of each M13 dye and 1 μL of the DNA template using a Bio-Rad thermocycler T100 (Bio-Rad, Pleasanton, CA, USA). PCR products were analyzed on an ABI 3500 genetic analyzer and sized against the LIZ500 internal standard (Applied Biosystems, Foster City, CA, USA). Allele calling was performed using Geneious software v. 9.1 (Kearse *et al.* 2012).

In order to estimate population differentiation and confirm species identity, two workers per nest were sequenced for a fragment of the cytochrome oxidase 1 marker (COI) using the *LF1* and *LR1* primer pair (Herbert et al. 2004; Smith *et al.* 2005). PCR products were purified using EXOSAP-it PCR purification kit (Applied Biosystems) and sequenced with the ABI BigDye Terminator v. 3.1 cycle sequencing kit (Applied Biosystems). Base calling and sequence pairing were performed using CodonCode Aligner (CodonCode Corporation, Dedham, MA, USA). Fifty-four new sequences were generated for our samples and combined with 27 additional sequences of *Melissotarsus* species obtained from GenBank (*M. insularis*, *M. weissi, M. beccarii* and *M. emeryi*; Smith *et al.* 2005).

### Population and colony structure

Allele frequencies, observed and expected heterozygosity, and *F*-statistics were assessed using FSTAT (Goudet 1995). Population and colony structure were determined using only worker genotypes. Every analysis was first performed at the entire population scale and successively applied for each locality. For each locality, genotypic frequencies were compared between every pair of nests using log-likelihood (G)-based tests of differentiation using GENEPOP ON THE WEB (Rousset 2008), to determine whether they belonged to the same colony. A Bonferroni correction was applied to account for multiple comparisons of all pairs. Significance was determined using a Fisher’s combined probability test. The genetic clustering of individuals within nests and populations was visualized by plotting individuals on a Principal Component Analysis (PCA) using *Adegenet R* package (Jombart 2008). The clustering of nests into distinct colonies was also assessed by Bayesian assignments of individuals into genetic clusters using STRUCTURE v. 2.3.4 (Pritchard et al. 2000). For each locality, the most likely number of genetic clusters (K) was estimated using simulations ran with values of K ranging from one to the total number of nests encountered within each dataset and repeated 10 times for each value of K. Each run included a 5 × 10^4^ burn in period followed by 1 × 10^5^ iterations of the MCMC. The most likely number of genetic clusters was evaluated using the ΔK method (Evanno et al. 2005) implemented in Structure Harvester v.0.6.8 (Earl & von Holdt 2012).

In addition, the phylogenetic relationship among mtDNA haplotypes was investigated using Maximum Likelihood implemented in the PhyML online web server (Guindon and Gascuel 2003). Nodal support was assessed by bootstrap resampling (1000 pseudoreplicates). Trees were visualized using FigTree v.1.3.1 (available at http://tree.bio.ed.ac.uk/software/figtree).

### Reproductive system and breeding strategies

For the colony fragments brought back to the lab, both alate and dealate queens were found while opening branches. The latter showed various degrees of gaster enlargement, with some being highly physogastric; ovaries and spermatheca were dissected in a subset of these queens. For all colonies, the presence of several reproductive queens was estimated using microsatellite markers, inferring whether all workers could be assigned to a single queen (carrying one of the two alleles of the mother queen at each microsatellite marker studied). The reproductive contribution of multiple queens was deduced when at least one worker per colony could not be unambiguously assigned to a single queen. The number of matings per queen was determined for each monogyne colony based on mother-offspring analyses using the maximum-likelihood method implemented in the software COLONY v.1.2 (Wang 2004). This analysis infers the number of males from the workers’ genotypes and assigns each worker to a given patriline.

For all colonies, relatedness coefficients (*r*) among nestmate workers were calculated using the program COANCESTRY v.1.0 (Wang 2011), according to the algorithm described by Queller & Goodnight (1989). Relatedness coefficients were weighted equally and standard errors (SE) were obtained by jackknifing over colonies. Relatedness coefficients were calculated separately for each locality to account for the strong genetic structure (and differences in allele frequencies) between populations (see Results).

## Results

### Population and colony structure

Overall, eight new haplotypes were found on the 622bp fragment of the COI mitochondrial marker for the 54 samples sequenced in this study. GenBank samples were included, although the species identifications may be ambiguous (*Melissotarsus* alpha taxonomy awaits a thorough revision). The overall dataset comprised 81 sequences for which 200 nucleotide positions were variable (175 parsimony-informative). Surprisingly, samples from different localities clustered into clearly separated clades (Figure S2). The Cederberg population clusters with the GenBank samples identified as *M. emeryi.* However, the other localities segregated away from *M. emeryi*, separated by one (*M. insularis,* for the localities of Eastern Cape and St Lucia) or even two species (*M. insularis* and *M. beccarii,* for the locality of uMkhuze). These findings suggests either that our sampling encompasses several undescribed species, or that previously described species represent geographical variants of a single species. In this study, we do not delimit different species; we hereafter refer them as different populations of *Melissotarsus beccarii,* as this species was first described in 1877 by Emery, and therefore has precedence over the others.

Nineteen microsatellite markers were successfully genotyped for up to eight workers per nest (mean ± SD = 7.49 ± 0.94; N = 262) and were found to be polymorphic with allele numbers ranging from 3 to 27 (mean ± SD = 11.26 ± 6.59). Strong genetic differentiation was found among nests of the overall population, with *F*_ST_ = 0.50 ± 0.15. Similar conclusions were reached within each locality, with significant population structure observed among nests within localities (*F*_ST_ = 0.29, 0.32, 0.32 and 0.27 for EC, MK, SL, and CE, respectively; *F_ST_* values for each pair of nests are given in Supplementary Information, Figure S3). Therefore, all but three pairs of nests were genetically different from each other (G-tests significant for each pair of nests; *P* < 0.001), even those located on the same host trees. The lack of differentiation between the three pairs of nests (EC1 and EC2; MK12 and MK3; SL10 and SL11), most likely results from their genetic similarity than their actual belonging to a common colony, as the nests from each pair were located on different trees, separated by hundreds of meters.

When the entire dataset from all locations was analyzed, STRUCTURE revealed the occurrence of at least three genetic clusters (most likely K = 3), roughly corresponding to the different localities sampled (Figure 2). When each locality was analyzed separately, STRUCTURE mostly segregated the different nests as distinct genetic clusters finding optimal K = 3 (out of 4 nests) for the EC population, K = 17 (out of 17) for the MK population, K = 8 (out of 9) for the SL population, and K = 3 (out of 3) for the CE population. The PCA analysis showed clear separation of localities and of nests within localities, while nestmate workers clustered together (Figure 3). Consistently with the genetic differentiation tests, the same three pairs of nests were also not separated from each other on both the STRUCTURE and PCA analyses. However, these nests are geographically distinct; their clustering therefore most likely stems from similar genetic diversity rather than belonging to a single colony. In general, the results at the overall population scale reflect the strong genetic differentiation observed between localities, which is consistent with the clear segregation observed on the mtDNA (Figure S2). The results at the locality scale indicate that nests mostly segregated from one another (Figure 4), confirming that every nest, even those sampled on the same host tree, represents its own separate colony. Interestingly, colonies *MK1* to *MK5* in the uMkhuze population were located on different stems of the same host tree; yet, the genetic differentiation between those colonies (pairwise *F_ST_* ± SD = 0.32 ± 0.07) was similar to the average *F_ST_* (= 0.33) observed within the uMkhuze population. However, the host tree *Strychnos madagascarensis* has an unusual growth pattern since new stems originate from the roots, and the stems never get tall (Suppl. Fig. S1). A clear differentiation between those colonies was expected as workers cannot walk between stems, hence colonies cannot merge together. However, similar differentiation was also found between colonies *MK7*, *MK8* and *MKF* (pairwise *F_ST_* ± SD = 0.34 ± 0.02; Figure S3) inhabiting a single branch of *S. madagascarensis*, and between colonies *SL1* to *SL3* (pairwise *F_ST_* ± SD = 0.23 ± 0.09) and colonies *SL4* and *SL5* (*F_ST_* = 0.43) inhabiting single ‘conventional’ trees in the St Lucia population (*F_ST_* = 0.32).

**Figure 2:**
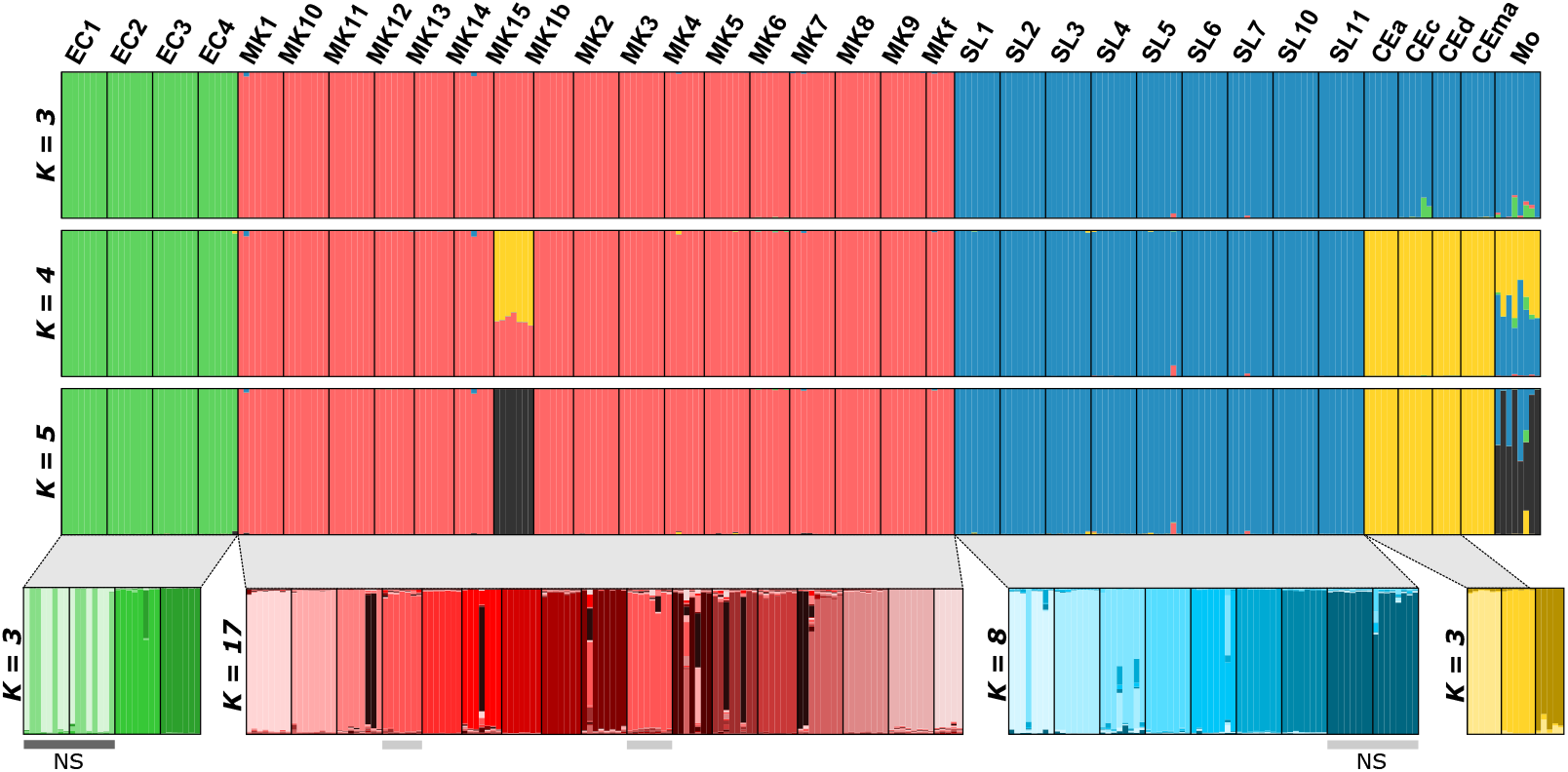
Graphical representation of STRUCTURE results determining the number of genetic groups in the overall dataset for different values of K. Each genetic group is characterized by a color; and each individual is represented by a vertical bar according to its probability of belonging to each group. Distinct simulations were subsequently run for the four populations, separately. In each population, gray bars indicate different colonies assigned to a single genetic group.

**Figure 3:**
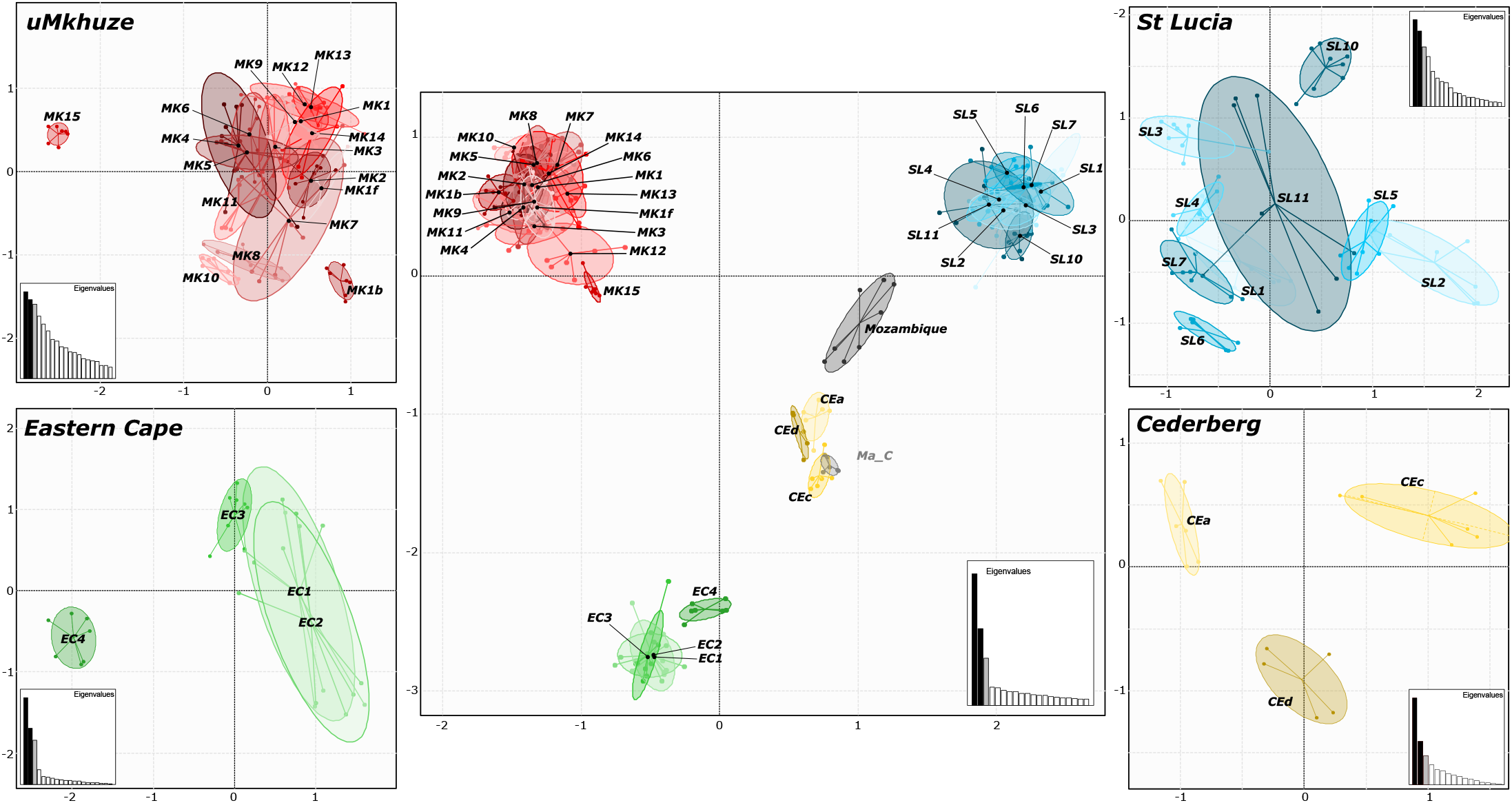
Clustering of nests in the overall sampling using principal component analysis of the microsatellite markers. Clustering analyses were subsequently run for each of the four populations of nests.

**Figure 4:**
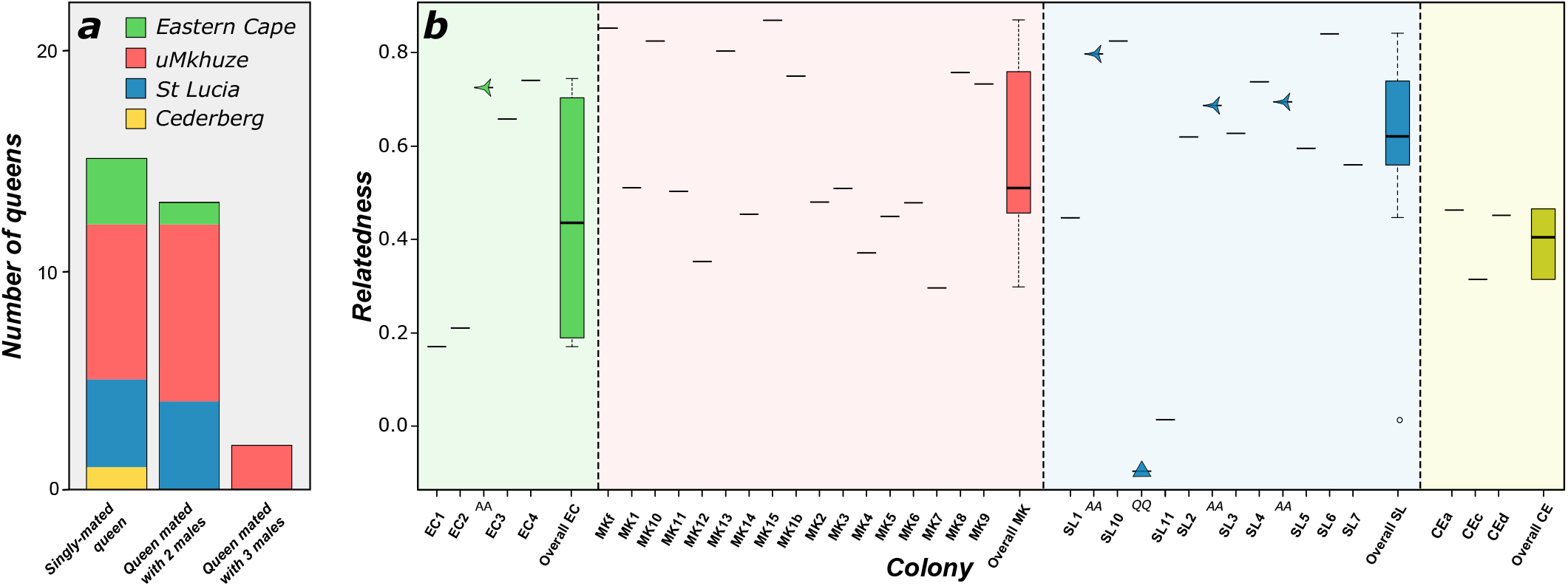
(*a*) Number of matings per queen for each monogyne colony in each population. (b) Relatedness values among nestmate workers for each colony. Arrows indicate relatedness values between alate queens (R_A-A_) and the triangle indicates relatedness value between queens in the *SL11* polygyne colony.

### Reproductive system and breeding strategies

A single reproductive queen was inferred genetically in all but one of the 34 colonies analyzed (*SL11* was polygyne). For monogyne colonies, mother-offspring inferences suggested that half of the queens were mated with a single male (16 out of the 33 queens analyzed), while 13 queens were mated with two males, and only two queens were mated with three males. No queen was found mated with more than three males (Figure 4a). Although a single reproducing queen was inferred in all colonies except *SL11* (see below), the genotypes of 4 out of 6 queens sampled are incompatible with being the mother of nestmate workers. Of these four colonies, the queens in *SL3* and *SL10* were related to the workers in their respective colonies (r_q-w_ ± SD = 0.18 ± 0.03 and 0.79 ± 0.0 for *SL3* and *SL10,* respectively), whereas the queens in *SL6* and *MK15* colonies were not (r_q-w_ ± SD = −0.26 ± 0.01 and −0.28 ± 0.0 for *SL6* and *MK15,* respectively). In contrast, the queens sampled in colonies *SL2* and *SL5* were the mothers of the workers from their respective colonies. Interestingly, different degrees of physogastry were observed among dealate queens (Figure 5). Ovarian dissections revealed that several non-physogastric queens are inseminated, but only some had active ovaries.

**Figure 5:**
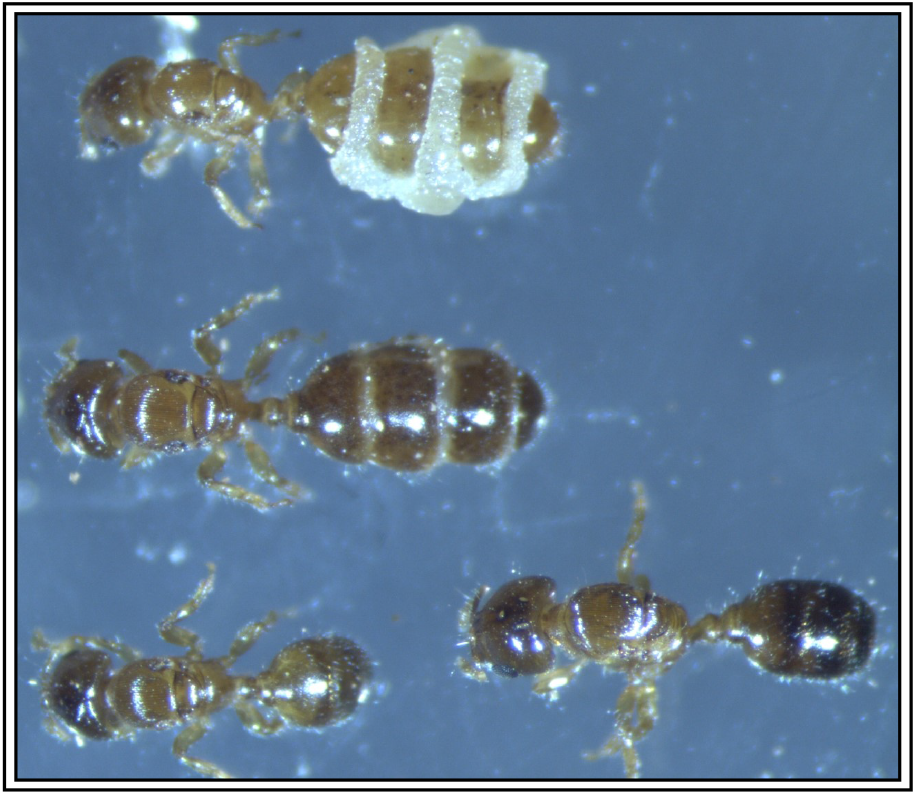
Dealate queens of *Melissotarsus* ant with different degrees of physogastry.

The occurrence of more than one reproductive queen was assessed in only one colony (*SL11* in St Lucia). Three mother queens were found while sampling a large branch in the tree housing colony *SL11*, but we cannot rule out the possibility that different adjacent colonies were pooled unintentionally. The genotypes of nominal *SL11* queens indicate that they are not related (r_q-q_ = −0.09). Notably, the increased genetic diversity resulting from the presence of multiple unrelated queens in *SL11* may explain the apparent lack of genetic differentiation with this colony in the STRUCTURE and PCA analyses. Interestingly, the inbreeding coefficient for the overall dataset was negative (*F*_IS_ ± SE = −0.32 ± 0.02), for the localities analyzed separately (*F*_IS_ = −0.23, −0.33, −0.34 and −0.31 for EC, MK, SL, and C, respectively), as well as for all colonies (except *SL11;* Figure S4). This absence of inbreeding reflects the non-independence of genotypes from a single family. This indicates that mother queens mated with unrelated males, while daughter queens and males do not stay and mate within the nest. Overall, within-colony relatedness was accordingly medium to high in most colonies, ranging from 0.17 (except r_w-w_ = 0.01 for *SL11*) to 0.87 (r_w-w_ ± SD = 0.55 ± 0.21; Figure 4b). Four nests had more than one alate queen. The genotypes of all alate queens analyzed (mean ± SD = 6 ± 2.1) was consistent with those of workers from the same colony, indicating that alate queens are produced by sexual reproduction like the workers, and all have the same mother. Consequently, the relatedness among alate queens within these four nests was also high, ranging from 0.69 to 0.87 (r_A-A_ ± SD = 0.72 ± 0.04; Figure 4b).

## Discussion

Our study provides valuable insights into the mating system and colony structure of wood-chewing *Melissotarsus* ants. Most colonies are headed by a single reproducing queen, mainly mated with a single male but occasionally mated with up to three males. In 4 out of 6 colonies for which a queen was sampled, we show that it did not produce the workers of the colony. This finding suggests relatively frequent queen turnover by related (n = 2) or unrelated queens (n = 2) that are re-accepted into already established colonies. We showed that colonies differ genetically from each other, even those located on different stems or branches of the same tree. We also highlighted that the four populations across South Africa exhibit strong genetic differentiation. A similar result was found with the mtDNA marker with each population representing a different clade, but they all exhibit a similar colony structure and breeding system. This absence of variation in mating strategies therefore suggests that the main evolutionary force reducing gene flow and enhancing speciation in this unorthodox genus is not related to differences in the breeding system.

The monodomous and monogynous colony structure observed in *Melissotarsus beccarii* in South Africa strongly differs from previous reports in Cameroon. In *M. beccarii* and *M. weissi*, numerous egg-producing physogastric queens were hypothesized to belong to the same colony because they were all found on the same tree, despite being located more than one meter away from each other (Mony et al. 2002). Low aggression between colonies was suggested to favor colonies merging as tunnels expand beneath the bark, resulting in a very populous, polygyne colony spanning an entire tree (Mony et al. 2007). In contrast with these observations, our results revealed that all nests sampled belong to distinct colonies, even those inhabiting the same branch. These findings therefore suggest that colonies maintain strict boundaries, casting doubt on the hypothesis that the multiple physogastric queens found on a tree (one meter apart) actually belong to the same colony. However, our study was performed on *Melissotarsus* in South Africa, and we therefore cannot rule out the possibility that distant populations or species of this genus exhibit different mating strategies.

Interestingly, in addition to the physogastric reproducing queen, numerous inseminated, but non-physogastric, queens were reported within Cameroon colonies of *M. beccarii* and *M. weissi* (Mony et al. 2002). In contrast with previous observations (Mony et al. 2002), our study reveals that some of them have active ovaries, and therefore seem to reproduce. However, our results indicate that all workers from all colonies studied can be assigned to a single mother (except that the mother queen can be physogastric or not), suggesting that these supplementary non-physogastric queens do not participate in worker production. In Cameroon, new alate queens are present within colonies and seem to swarm the year round (Mony et al. 2002). Our results showed that, within a colony, alate queens are produced by the same mother as the workers. We can therefore rule out the possibility that non-physogastric queens are opportunistic and produced reproductives, while the physogatric queen produces the numerous workers. Our sampling did not include enough males to conclude whether the non-physogastric queens with active ovaries may produce males within the colonies. Additionally, it is also possible that the current workers, through policing, cannibalize the eggs they produced (*i.e.,* whatever caste they would develop into).

The presence of these additional non-physogastric queens also questions whether they originated from the same physogastric mother, and whether they mate with their brothers in the nest to extend the colony lifespan after the mother queen dies. Mony et al. (2002) previously suggested that newly-inseminated queens are accepted by foreign colonies to perform worker-like tasks, and do not produce eggs until they have the chance to dominate their section of the colony (Mony et al. 2007). Our results partially support this hypothesis, as the queen sampled in 4 out of 6 colonies did not mother the workers of the colonies. For two of them, despite not being their mother, the new queens were related to the workers present in the colonies. The absence of inbreeding in any of the colonies sampled can rule out the possibility that these queens originated from the same mother *and* mate with their brothers (Trontti et al. 2005; Foitzik et al. 2011; Eyer et al. 2018b). Yet, we cannot exclude that these queens may originate from the same mother and mate with foreign males before reintegrating into their natal colony. This may happen through female-calling syndrome, whereby queens stand close to their nest entrance and release sex pheromones to attract neighboring males (Holldobler & Haskins 1977). Although this mating strategy leads to reduced gene flow compared to a nuptial flight of both sexes (Peeters & Aron 2017), it still prevents inbreeding, as males disperse from their colonies before mating (Bourke et al. 1988). For the two other colonies containing queens who were not the mothers of their nestmate workers, the new queens were not related to the workers present in the colonies. This result suggests queen turnover by unrelated foreign queens being accepted within the colonies, a finding previously suggested by Mony et al. (2002). Overall, these findings raise questions regarding the driving forces underlying queen turnover in this group, and prompt further investigation into the chemical and behavioral mechanisms determining queen acceptance, dominance and replacement in these species (Keller & Nonacs 1993; Koedam et al. 1997).

The mating strategy of *Melissotarsus* contrasts with other tree-living ant species, such as the devil garden’ ant *Myrmelachista schumanni,* the *Acacia* tree-living ant *Pseudomyrmex jerruginea,* or the *Cecropia* tree-living ants of the genus *Azteca.* In *My. schumanni*, workers selectively grow a ‘devil garden’ of their partner tree *Duroia hirsute* by attacking the saplings of other plants (Frederickson et al. 2005). This tree is mutualist with myrmecophytes, as it provides specific shelters for ants in their cavities (*i.e.,* domatia). Each garden of *My. schumanni* houses a single small supercolony, which contains up to three million workers freely moving across the hundreds of trees of the garden (Malé et al. 2020). Each garden may contain up to 15,000 queens but still maintains boundaries with different gardens and relatively high relatedness among nestmates through the reintroduction of daughter queens mated with related males (Malé et al. 2020). The ant species *P. jerruginea* and the swollen-thorn acacias *Acacia cornigera* are obligate mutualists, whereby the ant is dependent on the plant for food and dwelling, and the plant relies on the ant for protection against phytophagous insects and competing plants (Janzen 1966). In this ant species, a single queen establishes a colony digging in an unoccupied seedling or sucker shoot of a green thorn of the plant. As the colony grows, it further expands to the whole plant and often spreads out to neighboring acacias (up to 20), while the unique queen remains in the initial tree. As the colony expands, workers aggressively patrol the outside of the trees eliminating leaf herbivores, as well as any new foundresses over the next years. Interestingly, in other species of obligate acacia-ants (*e.g., P. belti*), newly mated queens are recruited as the colony growths. The large polygynous colonies can contain several million workers and several thousand queens. They can effectively patrol and monopolize several hundred of trees (Janzen 1966). In ants of the genus *Azteca,* multiple founding queens attempt to establish colonies, together or individually, on saplings of *Cecropia* tree (Longino 1989; Davidson et al. 1989; Yu & Davidson 1997). Each sapling may contain up to dozens of founding queens spread throughout the multiple internodes (Longino 1991). The queen or the association of several queens rear a first brood of workers without leaving the internode. Over time, the competition between queens restores monogyny within colonies, and the competition between colonies on each sapling ends up with a single colony occupying each *Cecropia* tree (Longino 1989).

The foraging strategy of *Melissotarsus* ants results in sharply contrasting colonization and cohabitation outcomes. Young trees are unsuitable for *Melissotarsus* ants because the bark is too thin to chew tunnels (likely to vary among tree genera). Only trees of a certain age are therefore suitable for colonization, while these trees also produce more branches that are appropriate over the years. In addition, the inability of workers to walk and patrol outside the host trees allow new foundresses to establish additional colonies in already habited trees, as long as they select ‘empty’ branches. Because chewing tunnels takes time, it is likely that empty branches are often available, and young queens can succeed to found new colonies. Overall, the foraging strategy of *Melissotarsus* ants allows multiple colonization events over many years on one same large tree, while the lack of encountering and direct competition between colonies allows several colonies to establish and cohabit on the same tree. Consequently, in contrast with other tree-living ant species, as well as with previous suggestions for this genus, *M. beccarii* colonies never reach colossal sizes (Mony et al. 2002); each tree rather contains a myriad of small *secluded* colonies of various ages.

Overall, these findings show that different phenologies of the plant host may select for distinct mating strategies and colony structures of the tree-living ant partner (Mayer et al. 2014). *Melissotarsus* species have been reported to inhabit at least 31 genera of host trees in 21 botanical families (Peeters et al. 2017), showing different growth and branching patterns, such as the multi-stemmed tree of the genus *Strychnos*. How the availability and the distribution of host trees shape colony structure, mating strategy and population genetics of the genus *Melissotarsus* clearly deserves further study.

## Acknowledgements

CP thanks Peter Hawkes, Peter Goodman, Robin Crewe and Bennie Bezuidenhout for logistical help in South Africa. Nigel Gericke and Peter Goodman helped with plant identification. Brigitte Church, iSimangaliso Wetland Park Authority and Ezemvelo KZN Wildlife, Funding for PAE and ELV was provided by the Texas A&M University Endowment in Urban Entomology.

## DATA ACCESSIBILITY

The microsatellite data reported in this study will be deposited in the Open Science Framework database upon acceptance, https://osf.io (DOI: XXXXX). The mtDNA sequences data generated in this study will be deposited in GenBank upon acceptance (accession numbers: XXXXX).

## AUTHOR CONTRIBUTIONS

CP designed the study. CP collected samples. PAE performed the genetic analyses and analyzed the data. PAE and CP wrote the paper with contributions of ELV.

## SUPPLEMENTARY MATERIAL

Additional material may be found in the online version of this article.

**Table S1**: Information on sampling location and number of individuals analyzed for each colony sampled.

**Table S2**: PCR multiplexing and number of alleles for each of the markers used in our study. This also includes the methods used to estimate detection of null alleles and linkage disequilibrium for the microsatellite markers analyses.

**Figure S1**: *Strychnos* and *Erythrina* trees that were sampled, showing the difference in growth patterns.

**Figure S2**: Maximum Likelihood tree of the mitochondrial COI marker. Each sample is colored according to its population of origin. Numbers indicate branch support through bootstrap values. Accession numbers and their described species are indicated for the samples from GenBank.

**Figure S3**: F_ST_ values based on microsatellite markers for each pair of nests.

**Figure S4**: Inbreeding coefficient (F_IS_ values) for each colony.

