## Supplementary material for "One tree, many colonies: colony structure, breeding system and colonization events of host trees in tunneling *Melissotarsus* ants": Table S1

**Supporting Information Table S1**

Information on sampling location and number of individuals analyzed for each colony sampled

| Colony | Population | Country | Notes on locality | Date | Same tree (T), branch (B) | Analyzed | | | |
|---|---|---|---|---|---|---|---|---|---|
| | | | | | | workers | Queens | Alate Q. | Males |
| *CE_a* | Cederberg_Western Cape | South Africa | Clanwilliam | (05-2017) | | 6 | | | |
| *CE_c* | Cederberg_Western Cape | South Africa | Clanwilliam | (05-2017) | | 6 | | | |
| *CE_d* | Cederberg_Western Cape | South Africa | Clanwilliam | (05-2017) | | 5 | | | |
| *Ma_C* | Cederberg_Western Cape | South Africa | Truitjieskraal | (06-2018) | | 5 | | | |
| *SL1* | St Lucia | South Africa | | (08-2019) | T1 | 8 | | | |
| *SL2* | St Lucia | South Africa | | (08-2019) | T1 | 8 | 1 | | |
| *SL3* | St Lucia | South Africa | | (08-2019) | T1 | 8 | 1 | 8 | |
| *SL4* | St Lucia | South Africa | | (08-2019) | T2 | 8 | | | |
| *SL5* | St Lucia | South Africa | | (08-2019) | T2 | 8 | 1 | 8 | |
| *SL6* | St Lucia | South Africa | | (08-2019) | | 8 | 1 | | |
| *SL7* | St Lucia | South Africa | | (08-2019) | | 8 | | 1 | |
| *SL10* | St Lucia | South Africa | | (08-2019) | | 8 | 1 | 5 | |
| *SL11* | St Lucia | South Africa | | (08-2019) | | 8 | 3 | | |
| *EC1* | East Cape | South Africa | near Cannon Rocks | (08-2019) | | 8 | | | |
| *EC2* | East Cape | South Africa | near Cannon Rocks | (08-2019) | | 8 | | | |
| *EC3* | East Cape | South Africa | near Cannon Rocks | (08-2019) | | 8 | | 3 | 3 |
| *EC4* | East Cape | South Africa | near Cannon Rocks | (08-2019) | | 8 | | | |
| *MK1b* | uMkhuze | South Africa | | (08-2019) | T3 | 7 | | | |
| *MK1* | uMkhuze | South Africa | | (08-2019) | T3 | 8 | | | |
| *MK2* | uMkhuze | South Africa | | (08-2019) | T3 | 8 | | | |
| *MK3* | uMkhuze | South Africa | | (08-2019) | T3 | 8 | | | |
| *MK4* | uMkhuze | South Africa | | (08-2019) | T3 | 8 | | | |
| *MK5* | uMkhuze | South Africa | | (08-2019) | T3 | 8 | | | |
| *MK6* | uMkhuze | South Africa | | (08-2019) | | 8 | | | |
| *MK7* | uMkhuze | South Africa | | (08-2019) | B1 | 8 | | | |
| *MK8* | uMkhuze | South Africa | | (08-2019) | B1 | 8 | | | |
| *MKF* | uMkhuze | South Africa | | (08-2019) | B1 | 5 | | | |
| *MK9* | uMkhuze | South Africa | | (08-2019) | | 8 | | | |
| *MK10* | uMkhuze | South Africa | | (08-2019) | | 8 | | | |
| *MK11* | uMkhuze | South Africa | | (08-2019) | | 8 | | | |
| *MK12* | uMkhuze | South Africa | | (08-2019) | | 7 | | | |
| *MK13* | uMkhuze | South Africa | | (08-2019) | | 7 | | | |
| *MK14* | uMkhuze | South Africa | | (08-2019) | | 7 | | | |
| *MK15* | uMkhuze | South Africa | | (08-2019) | | 7 | 1 | | |
| *Mo* | Northern Mozambique | Mozambique | | | | 8 | 4 | | 4 |
