## Supplementary material for "One tree, many colonies: colony structure, breeding system and colonization events of host trees in tunneling *Melissotarsus* ants": Table S2

**Supporting Information Table S2**

PCR multiplexing and number of alleles for each of the markers used in our study. This also includes the methods used to estimate detection of null alleles and linkage disequilibrium for the microsatellite markers analyses.

| Marker | PCR Mix | Dye | Range | Number of alleles |
|---|---|---|---|---|
| Ant859 | Mix1 | NED | 187-212 | 18 |
| Ant10878 | | NED | 314-318 | 4 |
| Ant7249 | | 6-FAM | 370-391 | 9 |
| Ant7680 | | 6-FAM | 299-320 | 13 |
| Ant5035 | | VIC | 296-311 | 8 |
| Ant1343 | | VIC | 249-264 | 12 |
| Ant575 | | PET | 254-281 | 12 |
| Ant8424 | Mix2 | NED | 212-225 | 6 |
| Ant3653 | | NED | 300-320 | 7 |
| Ant9218 | | 6-FAM | 397-434 | 13 |
| Ant2936 | | 6-FAM | 333-379 | 17 |
| Ant608 | | VIC | 195-215 | 5 |
| Ant3648 | | VIC | 318-334 | 11 |
| Ant3993 | | PET | 375-417 | 25 |
| Ant1368 | | PET | 303-343 | 21 |
| Ant11893 | Mix 3 | 6-FAM | 360-403 | 27 |
| Ant2341 | | VIC | 211-226 | 6 |
| Ant11315 | | VIC | 373-395 | 10 |
| Ant1387 | | PET | 180-184 | 3 |

Extracted DNA was amplified by PCR at 19 microsatellite loci (Butler et al. 2014) in 3 different mixes. Controls for genotyping errors due to null alleles were analysed following the Expectation Maximization algorithm of Dempster et al. (1977) implemented in the FREE NA software (Chapuis & Estoup 2007). Additional tests of heterozygote deficiency and estimation of linkage disequilibrium were performed in GENEPOP on the Web (Rousset 2000).

Butler IA, Siletti K, Oxley PR, Kronauer DJC. 2014. PLOS ONE 9(9): e107334

Chapuis M-P, Estoup A. 2007. Mol Biol Evol. 24: 621-631.

Rousset. 2000. J Evol Biol. 13: 58-62.

Dempster AP, Laird NM, Rubin DB. 1977. J R Stat Soc B. 39:1–38. 2.
