## Supplementary material for "One tree, many colonies: colony structure, breeding system and colonization events of host trees in tunneling *Melissotarsus* ants": Figure S1

**Conventional colony structure and mating strategy in the unorthodox tunneling ants of the genus *Melissotarsus***

*Supporting Information S1-continue*

Information on sampling location and number of individuals analyzed for each colony sampled

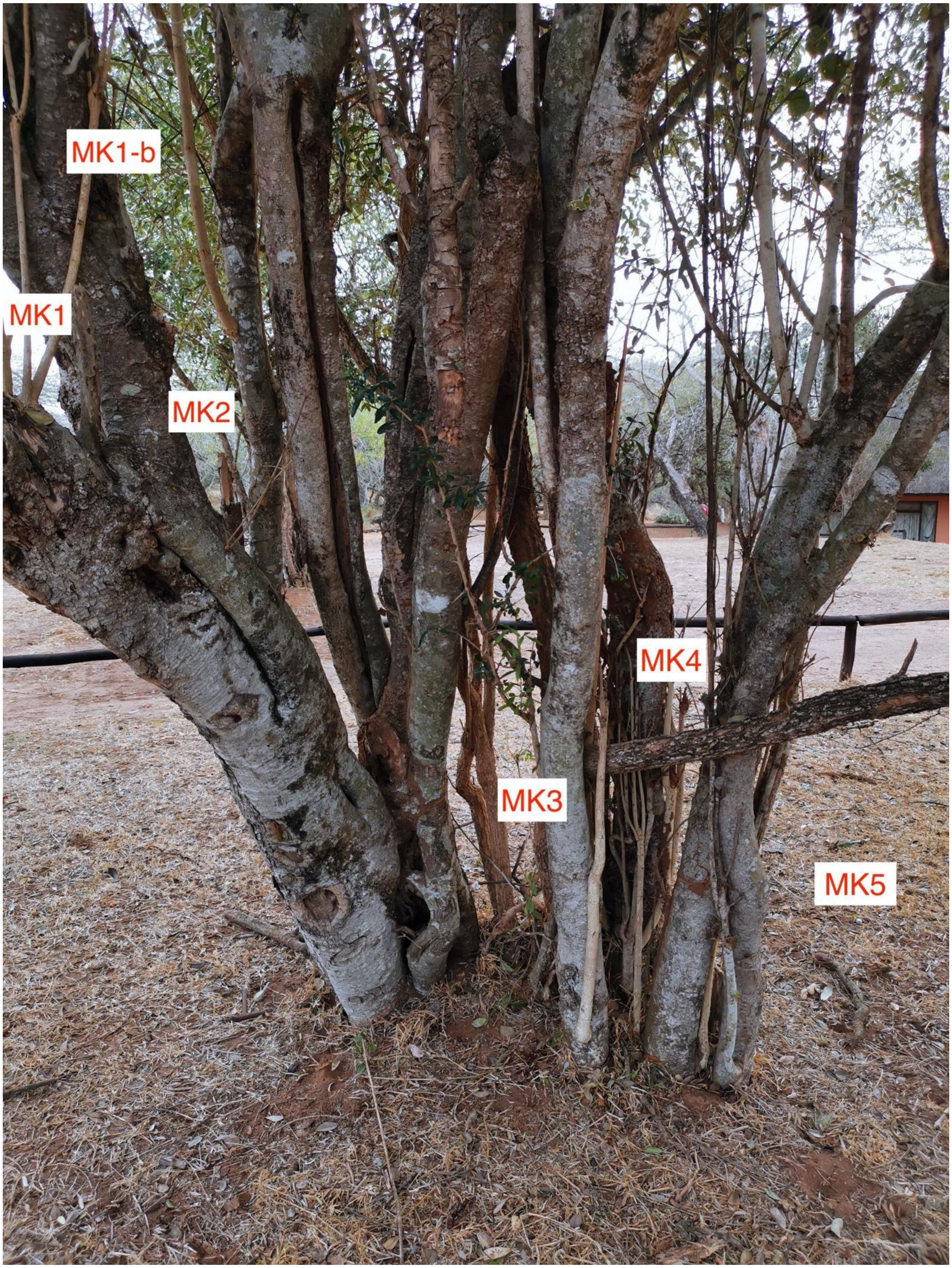


Colonies MK1 to MK5 located on a *Strychnos madagascariensis* tree in the uMkhuze population.

**Conventional colony structure and mating strategy in the unorthodox tunneling ants of the genus *Melissotarsus***

*Supporting Information S1-continue*

Information on sampling location and number of individuals analyzed for each colony sampled

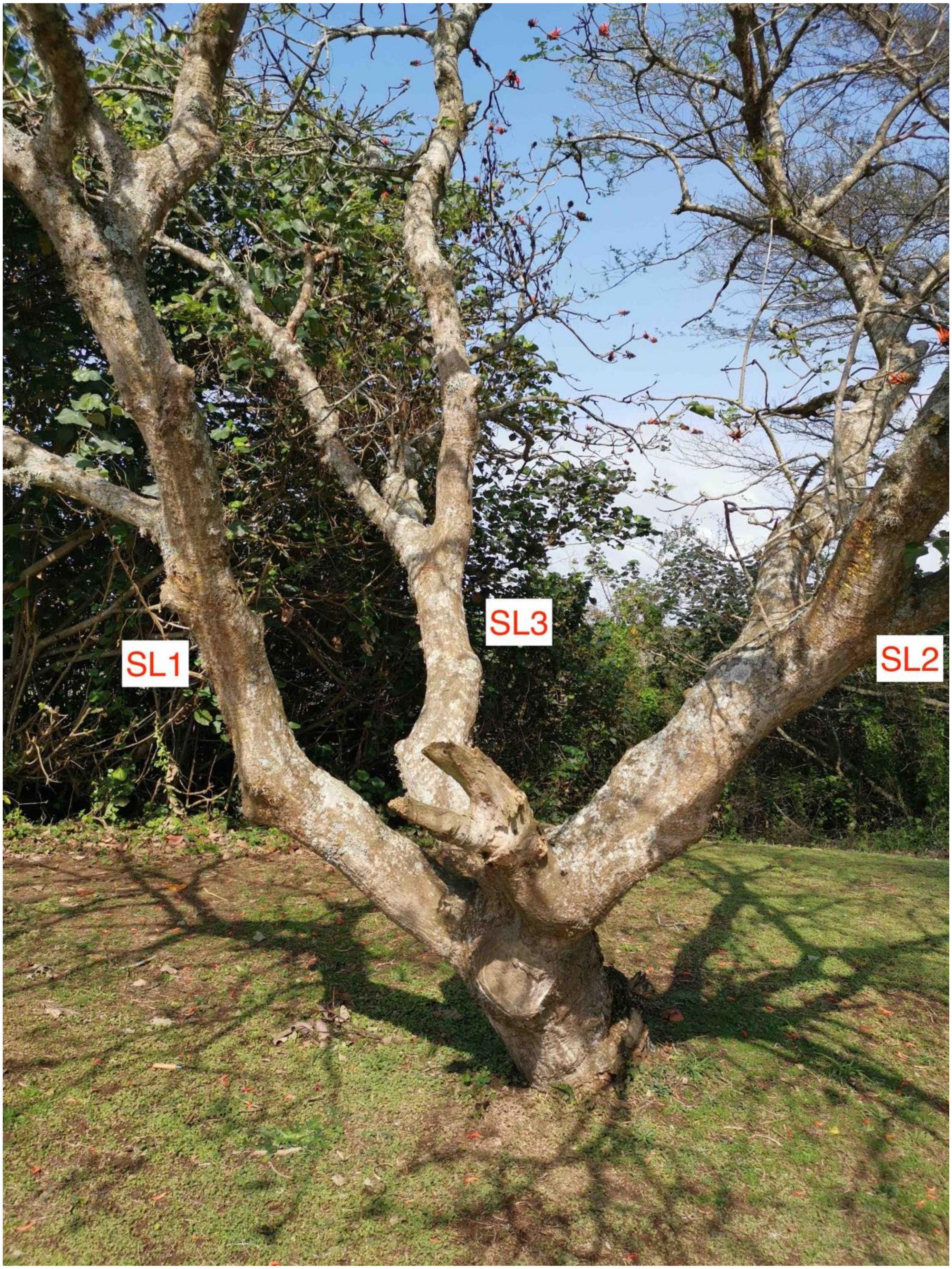


Colonies *SL1* to *SL3* located on a *Erythrina lysistemon* tree in the StLucia population.

**Conventional colony structure and mating strategy in the unorthodox tunneling ants of the genus *Melissotarsus***
*Supporting Information S1-continue*
Information on sampling location and number of individuals analyzed for each colony sampled

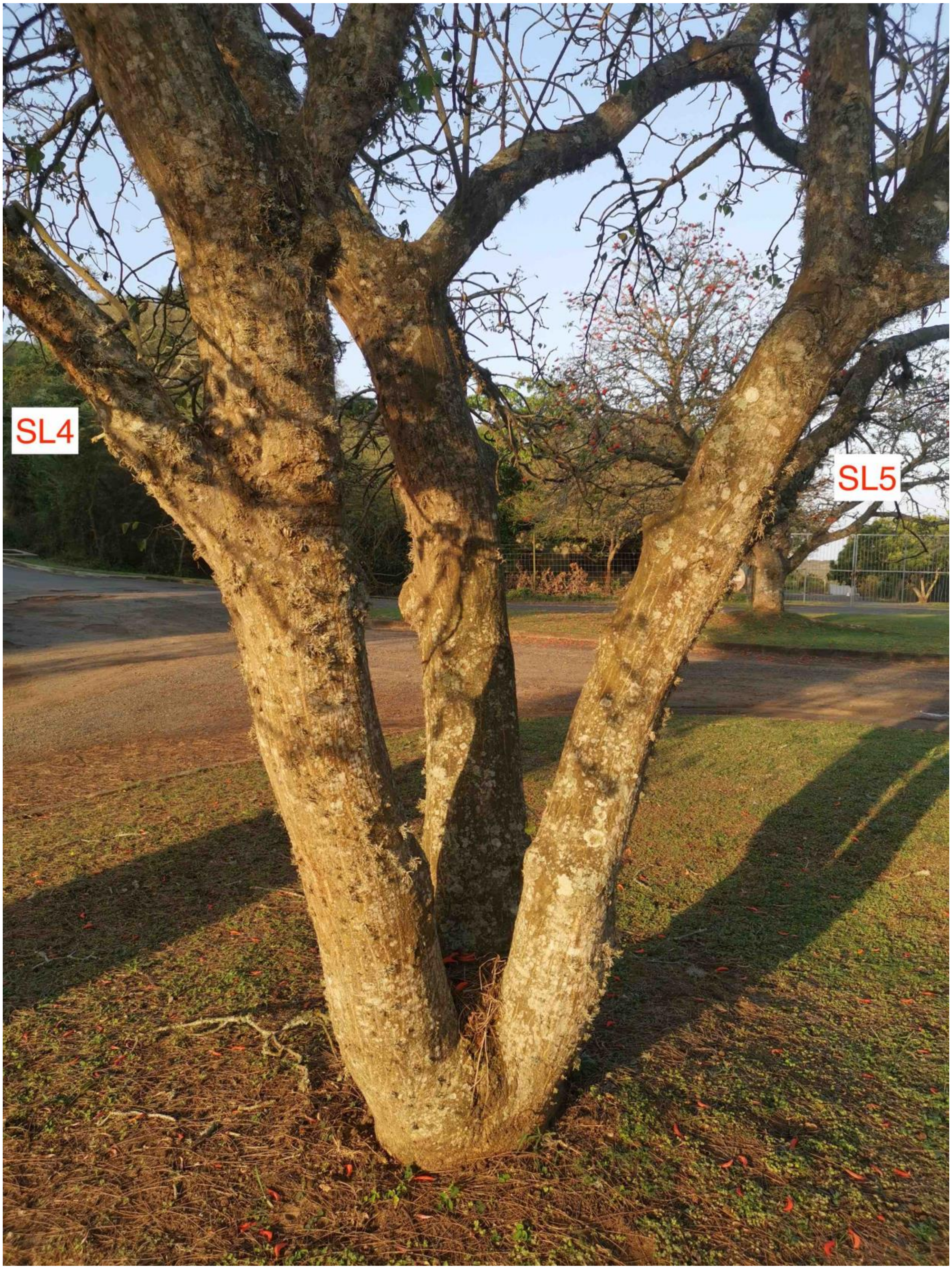


Colonies *SL4* to *SL5* located on a *Erythrina lysistemon* tree in the StLucia population.
