## Supplementary figures and images for "One tree, many colonies: colony structure, breeding system and colonization events of host trees in tunneling *Melissotarsus* ants"

### Figure S2

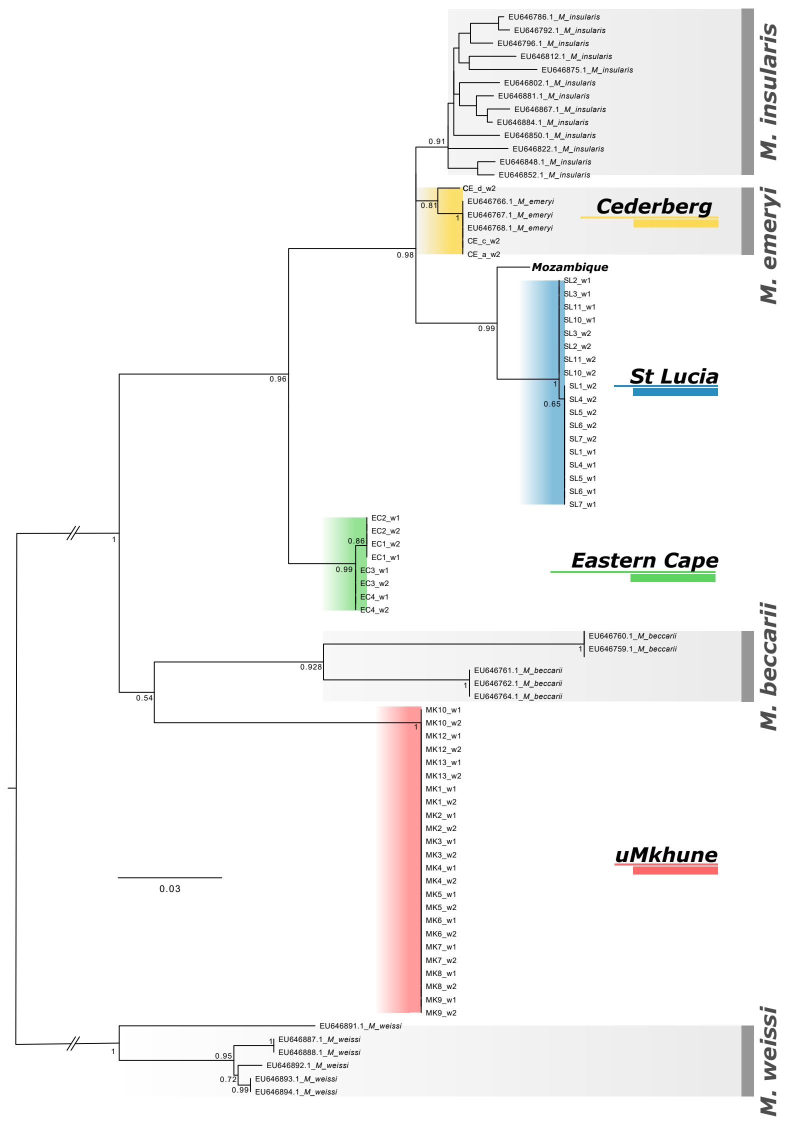

### Figure S4

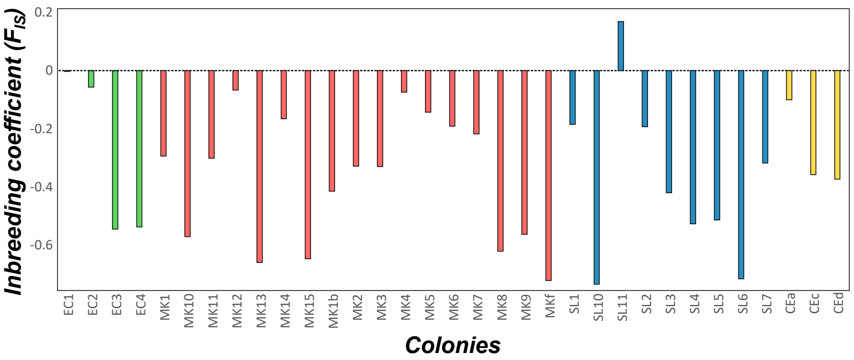
