## Supplementary material for "One tree, many colonies: colony structure, breeding system and colonization events of host trees in tunneling *Melissotarsus* ants": Figure S3

| | EC1 | EC2 | EC3 | EC4 | MK1 | MK1b | MK2 | MK3 | MK4 | MK5 | MK6 | MK7 | MK8 | MK9 | MK10 | MK11 | MK12 | MK13 | MK14 | MK15 | MKf | SL1 | SL2 | SL3 | SL4 | SL5 | SL6 | SL7 | SL10 | SL11 | CEa | CEc | CEd | CEma | Mo |
|---|---|---|---|---|---|---|---|---|---|---|---|---|---|---|---|---|---|---|---|---|---|---|---|---|---|---|---|---|---|---|---|---|---|---|---|
| EC1 | | -0.04 | 0.23 | 0.37 | 0.51 | 0.62 | 0.49 | 0.44 | 0.55 | 0.52 | 0.58 | 0.58 | 0.51 | 0.49 | 0.49 | 0.53 | 0.53 | 0.48 | 0.57 | 0.55 | 0.58 | 0.61 | 0.63 | 0.53 | 0.64 | 0.63 | 0.63 | 0.62 | 0.65 | 0.62 | 0.64 | 0.58 | 0.62 | 0.61 | 0.50 |
| EC2 | -0.04 | | 0.28 | 0.39 | 0.53 | 0.63 | 0.50 | 0.46 | 0.57 | 0.53 | 0.58 | 0.59 | 0.52 | 0.50 | 0.50 | 0.54 | 0.53 | 0.48 | 0.58 | 0.56 | 0.60 | 0.62 | 0.64 | 0.53 | 0.65 | 0.64 | 0.63 | 0.63 | 0.66 | 0.63 | 0.65 | 0.59 | 0.63 | 0.62 | 0.52 |
| EC3 | 0.23 | 0.28 | | 0.44 | 0.58 | 0.70 | 0.58 | 0.51 | 0.64 | 0.60 | 0.65 | 0.67 | 0.58 | 0.55 | 0.56 | 0.60 | 0.61 | 0.55 | 0.64 | 0.63 | 0.69 | 0.69 | 0.70 | 0.60 | 0.70 | 0.70 | 0.71 | 0.69 | 0.72 | 0.70 | 0.73 | 0.67 | 0.70 | 0.71 | 0.60 |
| EC4 | 0.37 | 0.39 | 0.44 | | 0.57 | 0.67 | 0.57 | 0.52 | 0.62 | 0.58 | 0.64 | 0.65 | 0.55 | 0.54 | 0.55 | 0.59 | 0.58 | 0.54 | 0.61 | 0.60 | 0.68 | 0.64 | 0.66 | 0.56 | 0.66 | 0.67 | 0.67 | 0.64 | 0.68 | 0.66 | 0.70 | 0.62 | 0.67 | 0.64 | 0.54 |
| MK1 | 0.51 | 0.53 | 0.58 | 0.57 | | 0.36 | 0.30 | 0.25 | 0.30 | 0.22 | 0.46 | 0.37 | 0.21 | 0.24 | 0.21 | 0.25 | 0.23 | 0.22 | 0.33 | 0.29 | 0.34 | 0.55 | 0.58 | 0.45 | 0.56 | 0.57 | 0.57 | 0.54 | 0.58 | 0.56 | 0.59 | 0.55 | 0.56 | 0.58 | 0.45 |
| MK1b | 0.62 | 0.63 | 0.70 | 0.67 | 0.36 | | 0.33 | 0.42 | 0.44 | 0.42 | 0.54 | 0.43 | 0.32 | 0.39 | 0.32 | 0.34 | 0.36 | 0.30 | 0.38 | 0.46 | 0.53 | 0.64 | 0.68 | 0.55 | 0.67 | 0.66 | 0.67 | 0.64 | 0.68 | 0.66 | 0.69 | 0.67 | 0.70 | 0.69 | 0.56 |
| MK2 | 0.49 | 0.50 | 0.58 | 0.57 | 0.30 | 0.33 | | 0.24 | 0.39 | 0.26 | 0.44 | 0.36 | 0.26 | 0.25 | 0.18 | 0.22 | 0.27 | 0.15 | 0.33 | 0.36 | 0.32 | 0.57 | 0.60 | 0.47 | 0.60 | 0.59 | 0.60 | 0.56 | 0.59 | 0.57 | 0.60 | 0.58 | 0.59 | 0.60 | 0.49 |
| MK3 | 0.44 | 0.46 | 0.51 | 0.52 | 0.25 | 0.42 | 0.24 | | 0.29 | 0.24 | 0.45 | 0.35 | 0.25 | 0.20 | 0.22 | 0.28 | 0.29 | 0.20 | 0.39 | 0.36 | 0.31 | 0.52 | 0.54 | 0.41 | 0.53 | 0.52 | 0.55 | 0.52 | 0.55 | 0.52 | 0.56 | 0.53 | 0.53 | 0.55 | 0.41 |
| MK4 | 0.55 | 0.57 | 0.64 | 0.62 | 0.30 | 0.44 | 0.39 | 0.29 | | 0.35 | 0.54 | 0.46 | 0.30 | 0.34 | 0.30 | 0.34 | 0.35 | 0.33 | 0.45 | 0.44 | 0.42 | 0.59 | 0.64 | 0.49 | 0.59 | 0.60 | 0.62 | 0.59 | 0.63 | 0.60 | 0.65 | 0.62 | 0.64 | 0.63 | 0.47 |
| MK5 | 0.52 | 0.53 | 0.60 | 0.58 | 0.22 | 0.42 | 0.26 | 0.24 | 0.35 | | 0.47 | 0.37 | 0.20 | 0.27 | 0.22 | 0.24 | 0.26 | 0.20 | 0.34 | 0.31 | 0.36 | 0.55 | 0.59 | 0.44 | 0.56 | 0.56 | 0.58 | 0.54 | 0.58 | 0.55 | 0.59 | 0.57 | 0.57 | 0.60 | 0.46 |
| MK6 | 0.58 | 0.58 | 0.65 | 0.64 | 0.46 | 0.54 | 0.44 | 0.45 | 0.54 | 0.47 | | 0.57 | 0.46 | 0.45 | 0.40 | 0.45 | 0.44 | 0.41 | 0.49 | 0.51 | 0.60 | 0.64 | 0.66 | 0.54 | 0.66 | 0.64 | 0.67 | 0.63 | 0.68 | 0.65 | 0.66 | 0.60 | 0.64 | 0.64 | 0.53 |
| MK7 | 0.58 | 0.59 | 0.67 | 0.65 | 0.37 | 0.43 | 0.36 | 0.35 | 0.46 | 0.37 | 0.57 | | 0.31 | 0.35 | 0.38 | 0.42 | 0.41 | 0.26 | 0.42 | 0.45 | 0.37 | 0.64 | 0.68 | 0.54 | 0.66 | 0.66 | 0.67 | 0.64 | 0.68 | 0.65 | 0.67 | 0.65 | 0.65 | 0.67 | 0.56 |
| MK8 | 0.51 | 0.52 | 0.58 | 0.55 | 0.21 | 0.32 | 0.26 | 0.25 | 0.30 | 0.20 | 0.46 | 0.31 | | 0.26 | 0.24 | 0.25 | 0.24 | 0.20 | 0.29 | 0.27 | 0.35 | 0.55 | 0.59 | 0.45 | 0.56 | 0.56 | 0.57 | 0.54 | 0.58 | 0.56 | 0.58 | 0.56 | 0.56 | 0.58 | 0.46 |
| MK9 | 0.49 | 0.50 | 0.55 | 0.54 | 0.24 | 0.39 | 0.25 | 0.20 | 0.34 | 0.27 | 0.45 | 0.35 | 0.26 | | 0.19 | 0.22 | 0.25 | 0.18 | 0.33 | 0.37 | 0.31 | 0.55 | 0.60 | 0.46 | 0.57 | 0.57 | 0.56 | 0.55 | 0.57 | 0.55 | 0.58 | 0.56 | 0.56 | 0.58 | 0.47 |
| MK10 | 0.49 | 0.50 | 0.56 | 0.55 | 0.21 | 0.32 | 0.18 | 0.22 | 0.30 | 0.22 | 0.40 | 0.38 | 0.24 | 0.19 | | 0.24 | 0.24 | 0.20 | 0.28 | 0.34 | 0.31 | 0.54 | 0.58 | 0.43 | 0.55 | 0.55 | 0.57 | 0.53 | 0.58 | 0.55 | 0.57 | 0.54 | 0.55 | 0.55 | 0.44 |
| MK11 | 0.53 | 0.54 | 0.60 | 0.59 | 0.25 | 0.34 | 0.22 | 0.28 | 0.34 | 0.24 | 0.45 | 0.42 | 0.25 | 0.22 | 0.24 | | 0.24 | 0.22 | 0.34 | 0.32 | 0.28 | 0.56 | 0.60 | 0.46 | 0.56 | 0.56 | 0.58 | 0.54 | 0.59 | 0.56 | 0.61 | 0.59 | 0.60 | 0.61 | 0.47 |
| MK12 | 0.53 | 0.53 | 0.61 | 0.58 | 0.23 | 0.36 | 0.27 | 0.29 | 0.35 | 0.26 | 0.44 | 0.41 | 0.24 | 0.25 | 0.24 | 0.24 | | 0.23 | 0.35 | 0.30 | 0.38 | 0.57 | 0.61 | 0.47 | 0.59 | 0.59 | 0.59 | 0.56 | 0.60 | 0.58 | 0.63 | 0.60 | 0.61 | 0.61 | 0.46 |
| MK13 | 0.48 | 0.48 | 0.55 | 0.54 | 0.22 | 0.30 | 0.15 | 0.20 | 0.33 | 0.20 | 0.41 | 0.26 | 0.20 | 0.18 | 0.20 | 0.22 | 0.23 | | 0.23 | 0.28 | 0.25 | 0.49 | 0.53 | 0.39 | 0.54 | 0.51 | 0.51 | 0.49 | 0.51 | 0.48 | 0.55 | 0.54 | 0.54 | 0.56 | 0.43 |
| MK14 | 0.57 | 0.58 | 0.64 | 0.61 | 0.33 | 0.38 | 0.33 | 0.39 | 0.45 | 0.34 | 0.49 | 0.42 | 0.29 | 0.33 | 0.28 | 0.34 | 0.35 | 0.23 | | 0.41 | 0.42 | 0.59 | 0.62 | 0.49 | 0.61 | 0.61 | 0.62 | 0.60 | 0.63 | 0.60 | 0.65 | 0.63 | 0.64 | 0.65 | 0.54 |
| MK15 | 0.55 | 0.56 | 0.63 | 0.60 | 0.29 | 0.46 | 0.36 | 0.36 | 0.44 | 0.31 | 0.51 | 0.45 | 0.27 | 0.37 | 0.34 | 0.32 | 0.30 | 0.28 | 0.41 | | 0.41 | 0.60 | 0.63 | 0.50 | 0.61 | 0.62 | 0.62 | 0.59 | 0.63 | 0.61 | 0.63 | 0.61 | 0.64 | 0.60 | 0.48 |
| MKf | 0.58 | 0.60 | 0.69 | 0.68 | 0.34 | 0.53 | 0.32 | 0.31 | 0.42 | 0.36 | 0.60 | 0.37 | 0.35 | 0.31 | 0.31 | 0.28 | 0.38 | 0.25 | 0.42 | 0.41 | | 0.66 | 0.68 | 0.53 | 0.69 | 0.66 | 0.67 | 0.64 | 0.71 | 0.66 | 0.74 | 0.70 | 0.69 | 0.71 | 0.54 |
| SL1 | 0.61 | 0.62 | 0.69 | 0.64 | 0.55 | 0.64 | 0.57 | 0.52 | 0.59 | 0.55 | 0.64 | 0.64 | 0.55 | 0.55 | 0.54 | 0.56 | 0.57 | 0.49 | 0.59 | 0.60 | 0.66 | | 0.36 | 0.15 | 0.36 | 0.30 | 0.28 | 0.25 | 0.22 | 0.19 | 0.63 | 0.60 | 0.63 | 0.62 | 0.40 |
| SL2 | 0.63 | 0.64 | 0.70 | 0.66 | 0.58 | 0.68 | 0.60 | 0.54 | 0.64 | 0.59 | 0.66 | 0.68 | 0.59 | 0.60 | 0.58 | 0.60 | 0.61 | 0.53 | 0.62 | 0.63 | 0.68 | 0.36 | | 0.23 | 0.46 | 0.40 | 0.47 | 0.36 | 0.47 | 0.39 | 0.69 | 0.65 | 0.70 | 0.66 | 0.42 |
| SL3 | 0.53 | 0.53 | 0.60 | 0.56 | 0.45 | 0.55 | 0.47 | 0.41 | 0.49 | 0.44 | 0.54 | 0.54 | 0.45 | 0.46 | 0.43 | 0.46 | 0.47 | 0.39 | 0.49 | 0.50 | 0.53 | 0.15 | 0.23 | | 0.24 | 0.22 | 0.24 | 0.20 | 0.22 | 0.14 | 0.52 | 0.49 | 0.51 | 0.50 | 0.30 |
| SL4 | 0.64 | 0.65 | 0.70 | 0.66 | 0.56 | 0.67 | 0.60 | 0.53 | 0.59 | 0.56 | 0.66 | 0.66 | 0.56 | 0.57 | 0.55 | 0.56 | 0.59 | 0.54 | 0.61 | 0.61 | 0.69 | 0.36 | 0.46 | 0.24 | | 0.43 | 0.46 | 0.29 | 0.45 | 0.40 | 0.67 | 0.64 | 0.67 | 0.63 | 0.41 |
| SL5 | 0.63 | 0.64 | 0.70 | 0.67 | 0.57 | 0.66 | 0.59 | 0.52 | 0.60 | 0.56 | 0.64 | 0.66 | 0.56 | 0.57 | 0.55 | 0.56 | 0.59 | 0.51 | 0.61 | 0.62 | 0.66 | 0.30 | 0.40 | 0.22 | 0.43 | | 0.36 | 0.38 | 0.40 | 0.29 | 0.64 | 0.61 | 0.63 | 0.61 | 0.45 |
| SL6 | 0.63 | 0.63 | 0.71 | 0.67 | 0.57 | 0.67 | 0.60 | 0.55 | 0.62 | 0.58 | 0.67 | 0.67 | 0.57 | 0.56 | 0.57 | 0.58 | 0.59 | 0.51 | 0.62 | 0.62 | 0.67 | 0.28 | 0.47 | 0.24 | 0.46 | 0.36 | | 0.36 | 0.42 | 0.31 | 0.69 | 0.66 | 0.70 | 0.67 | 0.45 |
| SL7 | 0.62 | 0.63 | 0.69 | 0.64 | 0.54 | 0.64 | 0.56 | 0.52 | 0.59 | 0.54 | 0.63 | 0.64 | 0.54 | 0.55 | 0.53 | 0.54 | 0.56 | 0.49 | 0.60 | 0.59 | 0.64 | 0.25 | 0.36 | 0.20 | 0.29 | 0.38 | 0.36 | | 0.36 | 0.30 | 0.65 | 0.63 | 0.66 | 0.62 | 0.39 |
| SL10 | 0.65 | 0.66 | 0.72 | 0.68 | 0.58 | 0.68 | 0.59 | 0.55 | 0.63 | 0.58 | 0.68 | 0.68 | 0.58 | 0.57 | 0.58 | 0.59 | 0.60 | 0.51 | 0.63 | 0.63 | 0.71 | 0.22 | 0.47 | 0.22 | 0.45 | 0.40 | 0.42 | 0.36 | | 0.18 | 0.68 | 0.65 | 0.69 | 0.66 | 0.44 |
| SL11 | 0.62 | 0.63 | 0.70 | 0.66 | 0.56 | 0.66 | 0.57 | 0.52 | 0.60 | 0.55 | 0.65 | 0.65 | 0.56 | 0.55 | 0.55 | 0.56 | 0.58 | 0.48 | 0.60 | 0.61 | 0.66 | 0.19 | 0.39 | 0.14 | 0.40 | 0.29 | 0.31 | 0.30 | 0.18 | | 0.64 | 0.62 | 0.65 | 0.63 | 0.44 |
| CEa | 0.64 | 0.65 | 0.73 | 0.70 | 0.59 | 0.69 | 0.60 | 0.56 | 0.65 | 0.59 | 0.66 | 0.67 | 0.58 | 0.58 | 0.57 | 0.61 | 0.63 | 0.55 | 0.65 | 0.63 | 0.74 | 0.63 | 0.69 | 0.52 | 0.67 | 0.64 | 0.69 | 0.65 | 0.68 | 0.64 | | 0.30 | 0.26 | 0.32 | 0.50 |
| CEc | 0.58 | 0.59 | 0.67 | 0.62 | 0.55 | 0.67 | 0.58 | 0.53 | 0.62 | 0.57 | 0.60 | 0.65 | 0.56 | 0.56 | 0.54 | 0.59 | 0.60 | 0.54 | 0.63 | 0.61 | 0.70 | 0.60 | 0.65 | 0.49 | 0.64 | 0.61 | 0.66 | 0.63 | 0.65 | 0.62 | 0.30 | | 0.23 | 0.01 | 0.47 |
| CEd | 0.62 | 0.63 | 0.70 | 0.67 | 0.56 | 0.70 | 0.59 | 0.53 | 0.64 | 0.57 | 0.64 | 0.65 | 0.56 | 0.56 | 0.55 | 0.60 | 0.61 | 0.54 | 0.64 | 0.64 | 0.69 | 0.63 | 0.70 | 0.51 | 0.67 | 0.63 | 0.70 | 0.66 | 0.69 | 0.65 | 0.26 | 0.23 | | 0.21 | 0.45 |
| CE_ma | 0.61 | 0.62 | 0.71 | 0.64 | 0.58 | 0.69 | 0.60 | 0.55 | 0.63 | 0.60 | 0.64 | 0.67 | 0.58 | 0.58 | 0.55 | 0.61 | 0.61 | 0.56 | 0.65 | 0.60 | 0.71 | 0.62 | 0.66 | 0.50 | 0.63 | 0.61 | 0.67 | 0.62 | 0.66 | 0.63 | 0.32 | 0.01 | 0.21 | | 0.47 |
| Mo | 0.50 | 0.52 | 0.60 | 0.54 | 0.45 | 0.56 | 0.49 | 0.41 | 0.47 | 0.46 | 0.53 | 0.56 | 0.46 | 0.47 | 0.44 | 0.47 | 0.46 | 0.43 | 0.54 | 0.48 | 0.54 | 0.40 | 0.42 | 0.30 | 0.41 | 0.45 | 0.45 | 0.39 | 0.44 | 0.44 | 0.50 | 0.47 | 0.45 | 0.47 | |
